# Tauopathy and alcohol consumption interact to alter locus coeruleus excitatory transmission and excitability in male and female mice

**DOI:** 10.1101/2022.07.22.501151

**Authors:** Anthony M. Downs, Christina M Catavero, Michael R. Kasten, Zoé A. McElligott

## Abstract

Alcohol use disorder is a major public health concern in the United States. Recent work has suggested a link between chronic alcohol consumption and the development of tauopathy disorders, such as Alzheimer’s Disease and Frontal-Temporal Dementia. However, relatively little work has investigated changes in neural circuitry involved in both tauopathy disorders and alcohol use disorder. The locus coeruleus (LC) is the major noradrenergic nuclei in the brain and is one of the earliest sites to be affected by tau lesions. The LC is also implicated in the rewarding effects of ethanol and alcohol withdrawal. In this study we assessed effects of long-term ethanol consumption and tauopathy on the physiology of LC neurons. Male and female P301S mice, a humanized transgenic mouse model of tauopathy, underwent 16 weeks of intermittent access to 20% ethanol from 3 to 7 months of age. We observed higher total alcohol consumption in female mice regardless of genotype. Male P301S mice consumed more ethanol and had a greater preference for ethanol than WT males. At the end of the drinking study, LC function was assessed using *ex vivo* whole cell electrophysiology. We found significant changes in excitatory inputs to the LC due to both ethanol and genotype., We found significantly increased excitability of the LC due to ethanol with greater effects in female P301S mice than WT. Our study identifies significant changes in the LC due to interactions between tauopathy and long-term ethanol use. These findings could have important implications regarding LC activity and changes in behavior due to both ethanol and tauopathy related dementia.

## INTRODUCTION

Over the past several decades, there has been an increase in the incidence of Alzheimer’s Disease (AD) along with an increase in alcohol consumption in adults 50 years and older (Breslow et al., 2017; Grucza et al., 2018; Masters et al., 2015; Mayeux and Stern, 2012; Wilson et al., 2014). Heavy alcohol consumption throughout life may enhance the risk for developing dementia from tauopathy disorders, including AD and frontotemporal dementia (FTD) (Heymann et al., 2016; Peng et al., 2020; Rehm et al., 2019; Tyas, 2001; Wang et al., 2021). Mouse models of AD have also demonstrated increased AD-associated pathology due to alcohol consumption (Barnett et al., 2022; Hoffman et al., 2019; Tucker et al., 2022). While there appears to be a clear linkage between alcohol use, the ageing brain, and tauopathies including FTD and AD, there has been relatively little work examining the interaction of tau and alcohol in specific neural circuits.

The locus coeruleus (LC) is a major site of early tauopathy and is thought to be critical for the pathogenesis of AD and FTD (Beardmore et al., 2021; Ohm et al., 2020). The LC is a dense collection of noradrenergic neurons located in the pons and innervates many structures throughout the neuroaxis (Poe et al., 2020). Due to this wide innervation pattern, the LC is a neuronal hub important for a variety of functions including arousal, sleep-wake cycles, stress responses, memory and leaning, and cognition (Aston-Jones and Cohen, 2005; Harley, 1987; Kayama and Koyama, 2003; Mason, 1981). The LC is one of the first sites of neuropathology in AD and roughly 70% of LC neurons are lost by terminal stages of AD in humans (Andrés-Benito et al., 2017; Bondareff et al., 1982; Braak and Braak, 1991; Rüb et al., 2001; Zweig et al., 1988). Loss of LC neurons and noradrenergic dysfunction is associated with cognitive decline in AD and restoring LC function has been suggested as a therapeutic target for AD (Ciampa et al., 2022; Jacobs et al., 2021; Rorabaugh et al., 2017). Animal models also support the involvement of the LC in the development of dementia. Injection of synthetic tau fibrils in the *P301S MAPT* (PS19) transgenic mouse, results in the spread of tau according to LC afferent projections (Iba et al., 2015). Further, ablation of the LC from *P301S MAPT* transgenic mice exacerbates cognitive deficits and neuropathology (Chalermpalanupap et al., 2018). Accordingly, tau related dysfunction in the LC is well positioned to disrupt a variety of behavioral and cognitive processes.

The LC is also a critical nucleus for alcohol use disorder (AUD) and the rewarding effects of ethanol (Cannady et al., 2018; Downs and McElligott, 2022; Harrison et al., 2017; Vazey et al., 2018). Injection or binge consumption of alcohol induces cFos expression in the LC (Burnham and Thiele, 2017; Thiele et al., 2000). Global deletion of *DBH* (Dopamine β-hydroxylase), reduces alcohol consumption and preference in mice (Weinshenker et al., 2000). α1 and β andrenoreceptor (AR) antagonists and α2 AR agonists have also been demonstrated to reduce alcohol cravings and the negative affective symptoms of alcohol withdrawal in several clinical trials (Bailly et al., 1992; Gottlieb et al., 1994; Haass-Koffler et al., 2017; Horwitz et al., 1989; Keaney et al., 2001; Milivojevic et al., 2020; Simpson et al., 2015; Simpson et al., 2018; Sinha et al., 2021). Previous studies have demonstrated increased LC neuron excitability following abstinence from alcohol in rats and mice, due to increased excitatory amino acid receptors and alterations in corticotrophin-releasing factor innervation from the central nucleus of the amygdala (Engberg and Hajós, 1992; Funk et al., 2006; Gilpin et al., 2015; Koob, 2015). Together, this suggests that chronic alcohol consumption and tau burden may interact to potentiate LC dysfunction.

To investigate the interaction between alcohol consumption and tauopathy on LC neuron function, we exposed *P301S MAPT* transgenic mice and wild-type (WT) littermates to 16-weeks of intermittent access (IA) to ethanol. After 16-weeks of IA and when the mice were 7 months of age, we assessed LC physiology using *ex vivo* whole-cell patch-clamp electrophysiology. We found that female WT and P301S mice drank significantly more ethanol than males. We found significant decreases in both frequency and amplitude of spontaneous excitatory postsynaptic currents (sEPSCs) in male P301S mice. However, female mice showed changes in sEPSC frequency and kinetics due to both P301S genotype and IA to ethanol. We did not find any significant changes in the spontaneous action potential (AP) firing rate of LC neurons due to either P301S genotype or IA to ethanol in either males or females. However, we did see a significant increase in ethanol-induced excitability of LC neurons in both P301S and WT mice of both sexes. Taken together, these results demonstrate significant plasticity of LC neurons in response to alcohol and tau pathology, many of which are also modified by sex.

## MATERIALS AND METHODS

### Animals

Adult male and female P301S (Jackson Labs #008160) mice on a C57BL/6J background and wildtype littermate controls were bred in house. All animals in the study were 12-13 weeks old at the onset of drinking studies. Animals were singly housed and maintained on a reverse 12-hour light cycle with lights off from7:00am-7:00pm. Animals had *ad libitum* access to food and water. All animal procedures were performed in accordance with the regulations of the University of North Carolina at Chapel Hill’s institutional animal care and use committee. The animals used in this study are a subset of the animals reported in (Catavero et al., 2022). These animals were included in the drinking metrics, but not the behavioral experiments.

### Ethanol consumption paradigm

Intermittent access (IA) to ethanol was performed in home cages as previously described (Hwa et al., 2011). Briefly, experimental mice were housed singly and provided access to an RHM 3000 diet (Prolab) one week prior to ethanol drinking. Mice were then provided access to a bottle of ethanol and water in their home cage on Monday, Wednesday, and Friday for 24-hour periods with bottles changed in the A.M. On other days, they had access to two bottles of water. Ethanol and water bottles were rotated to prevent association of ethanol with a particular side of the cage. Control mice had access to two bottles of water every day. Mice underwent the intermittent access paradigm for sixteen weeks beginning at 12-13 weeks of age. Experimental mice had access to 3%, 6%, 10% (v/v) ethanol (unsweetened) for weeks 1-3 of the experiment, and access to 20% (v/v) ethanol (unsweetened) for weeks 4-16 of the experiment. Our design provides 4 groups for each sex: MWW-<u>m</u>ale <u>w</u>ildtype <u>w</u>ater, MWE-<u>m</u>ale <u>w</u>ildtype <u>e</u>thanol, MPW-<u>m</u>ale <u>P</u>301S <u>w</u>ater, MPE-<u>m</u>ale <u>P</u>301S <u>e</u>thanol, FWW-<u>f</u>emale <u>w</u>ildtype <u>w</u>ater, FWE-<u>f</u>emale <u>w</u>ildtype <u>e</u>thanol, FPW-<u>f</u>emale <u>P</u>301S <u>w</u>ater, and FPE-<u>f</u>emale <u>P</u>301S <u>e</u>thanol. Mice then underwent a 72-hour forced abstinence period from alcohol before being euthanized for electrophysiological studies at approximately 7 months of age (Fig 1A). All recordings were performed within 1 week of the end of the 72-hour forced abstinence period. 7 months of age was chosen because there is evidence of significant tau accumulation at this time point, without the significant mortality seen at later time points (Yoshiyama et al., 2007).

**Figure 1.**
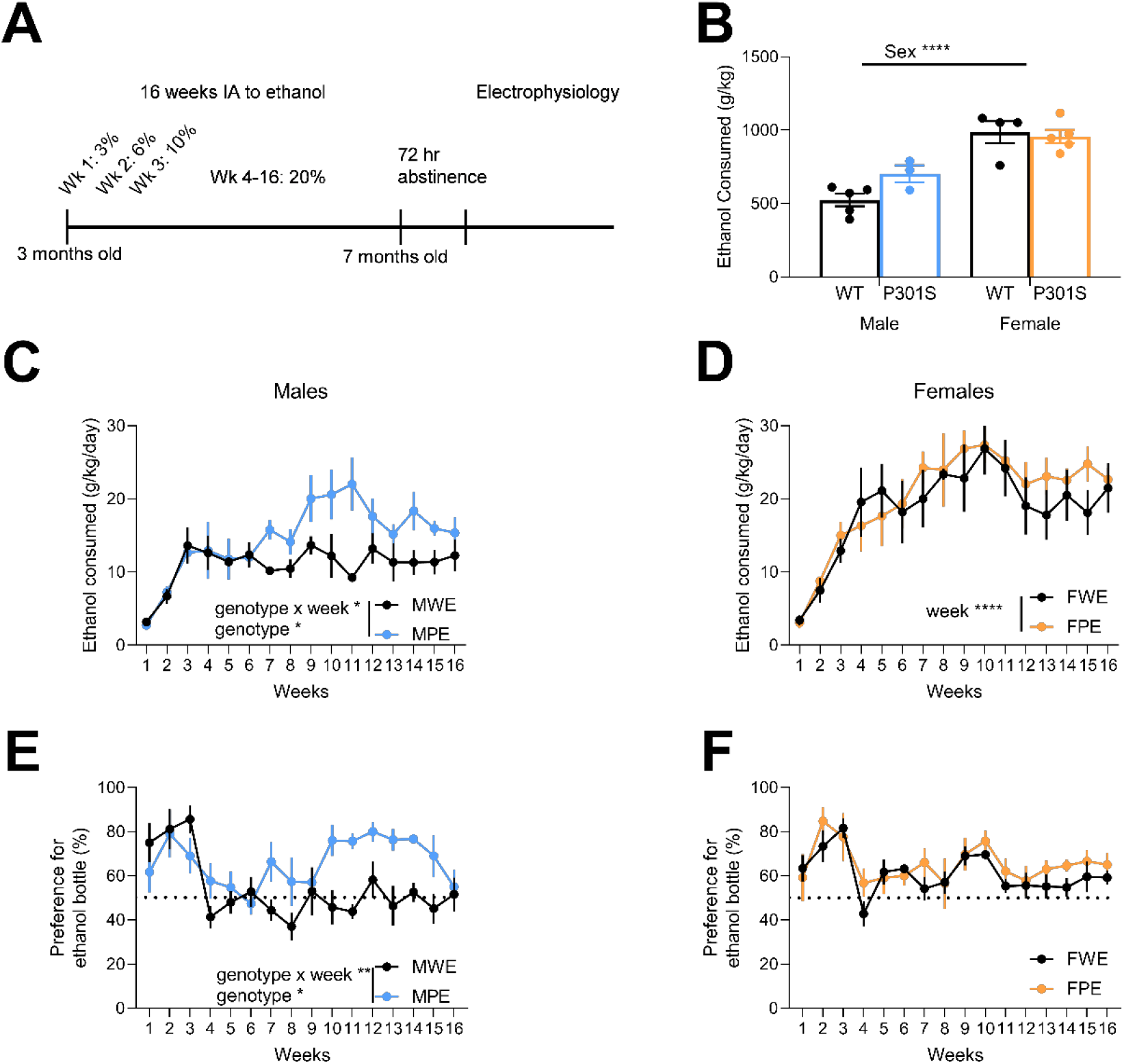
Ethanol consumption in intermittent access paradigm. (A) Schematic of experimental design (B) Total ethanol consumption over 16 weeks. Females consumed more ethanol than males regardless of genotype. (C) Weekly ethanol consumption in males. Male mice had escalated ethanol consumption across 16 weeks, and MPE mice consumed more alcohol than MWE mice over later weeks. (D) Weekly ethanol consumption in females. Female mice had escalated ethanol consumption across 16 weeks. (E) Weekly preference for ethanol bottle in males. MPE mice developed a greater preference for the ethanol bottle over later weeks. (F) Weekly preference for ethanol bottles in females. There was no difference in preference between FWE and FPE mice. Data expressed as mean ± S.E.M. The following sample sizes were used: MWW n = 4 mice, MWE n = 5 mice, MPW n = 5 mice, MPE n =3 mice, FWW n = 4 mice, FWE n = 4 mice, FPW n = 4 mice, FPE n = 5 mice. * p ≤ 0.05, ** p ≤ 0.01, **** p ≤ 0.0001.

### Brain Slice Preparation

We prepared brain slices for whole-cell electrophysiology as previously described (Luster et al., 2020). Briefly, mice were deeply anesthetized with isoflurane, decapitated, and brains were removed and placed into ice-cold sucrose aCSF [in mM: 194 sucrose, 20 NaCl, 4.4 KCl, 2 CaCl_2_, 1.2 NaH_2_PO_4_. 10 glucose, 26 NaHCO_3_] oxygenated with 95% O_2_ 5% CO_2_ for slicing. Brains were sliced coronally at 200 μm using a Leica VT1000 vibratome (Germany). Brain slices were then incubated in oxygenated aCSF [in mM: 124 NaCl, 4.4 KCl, 2 CaCl_2_, 1.2 MgSO_4_. 1 NaH_2_PO_4_, 10 glucose, 26 NaHCO_3_] held at 32 °C for at least 45 minutes prior to recording. Slices were then transferred to a recording chamber and perfused with oxygenated aCSF at 28-30 °C at a constant rate of 2 mL/min.

### Whole-cell recordings

Recording electrodes (2-4 MΩ) were pulled on a P-97 Micropipette Puller (Sutter Instruments). All recordings were conducted with potassium-gluconate intercellular recording solution [in mM: 135 gluconic acid potassium, 5 NaCl, 2 MgCl_2_, 10 HEPES, 0.6 EGTA, 4 Na_2_ATP, 0.4 Na_2_GTP). All signals were acquired using an Axon Multiclamp 700B (Molecular Devices Sunnyvale, CA). Input resistance, holding current, and access resistance were continuously monitored throughout the experiment. Cells in which access resistance changed greater than 20% were excluded from analysis. 2-4 cells were recorded from each animal for each set of experiments.

LC neurons were identified by their anatomical location in the slice, morphology, absence of Ih currents, spontaneous action potentials, and linear response to hyperpolarizing currents (Paladini et al., 2007; Williams et al., 1984). Spontaneous glutamate-mediated excitatory postsynaptic currents (sEPSCs) were acquired in voltage clamp at a −80 mV holding potential. Spontaneous action potentials were acquired in current-clamp mode with no current injected (I=0). Excitability studies were performed in current clamp mode with current injected to hold cells to a common membrane potential of −75 mV to account for inter-cell variability. Variation in excitability was then assessed with negative current injection (−100 - 0 pA at 20 pA intervals) to assess hyperpolarizing sag, and with positive current injection (0 – 400 pA at 20 pA intervals) to assess the number of action potentials following each current step. The same cells were used for both current clamp and voltage clamp experiments.

### Data analysis

All electrophysiological data was analyzed using pClamp 10.6 software (Molecular Devices) and Minianalysis (Synaptosoft). All data were analyzed using a two/three way regular or repeated measures ANOVA depending on the number of variables assessed in a given experiment. Male and female data were analyzed separately in all electrophysiological experiments due to significant differences in ethanol consumption. Significant main or interaction effects were subsequently analyzed using Tukey’s multiple comparisons test to control for multiple comparisons. All data are expressed as mean ± SEM. P values ≤ 0.05 were considered statistically significant. All statistical tests were performed using Graphpad Prism 9.30 (La Jolla, CA, USA).

## RESULTS

### Alcohol Consumption

Because the IA paradigm used in this study is a voluntary consumption model, we first assessed total ethanol consumption during the 16 weeks of IA. Both WT and P301S female mice drank significantly more total ethanol than male mice of either genotype (main effect of sex, F_1,13_ = 40.21, p < 0.0001, Tukey’s multiple comparisons test, MWE vs. FWE p = 0.0002, MPE vs FPE p = 0.406) (Fig 1B). There was no difference in total ethanol consumption over the duration of the experiment due to genotype (main effect of genotype, F_1,13_ = 1.718, p = 0.21). Both male and female mice escalated their ethanol intake over progressive weeks, which is consistent with previous studies using the IA model (main effect of week, F_3.46,20.73_ = 7.32, p = 0.001, Fig 1C & D, (Hwa et al., 2011). However, MPE mice drank more ethanol as the weeks progressed than MWE mice (significant interaction effect, F_15,90_ = 2.15, p = 0.014; main effect of genotype, F_1,6_ = 6.21, p = 0.047). We saw no difference in ethanol consumption between FWE and FPE mice (main effect of genotype, F_1,7_ = 0.049, p = 0.51) (Fig 1D). To further examine differences in ethanol consumption in males, we examined preference for the ethanol bottle across weeks. We found that MPE mice had a greater preference for the ethanol bottle relative to MPW mice (week x genotype interaction effect, F_15,90_ = 2.37, p = 0.006; main effect of genotype, F_1,6_ = 6.81, p = 0.04) (Fig 1E). We did not observe any differences in ethanol preference between FWE and FPE animals (main effect of genotype, F_1,8_ = 1.84, p = 0.21) (Fig 1F). Due to the differences in total alcohol consumption between male and female mice, males and females were analyzed separately for the subsequent electrophysiological studies.

### Electrophysiological properties of LC neurons

Because the LC is known to be modified by both ethanol consumption and tau pathology (Beardmore et al., 2021; Downs and McElligott, 2022), we next assessed baseline whole cell resistance and capacitance of LC neuron in voltage clamp mode using steps from −70 to −80 mV. We observed an increase in whole cell resistance in both WT and P301S male mice in response to ethanol drinking (main effect of ethanol, F_1,31_ = 5.98, p = 0.023) (Fig 2A). We also observed a decrease in cell capacitance in male P301S mice versus WT animals (main effect of genotype, F_1,30_ = 4.36, p = 0.046) (Fig 2B). However, we did not observe any changes in whole cell resistance in female mice regardless of genotype (Fig 2C). We observed a significant decrease in capacitance due to ethanol in both WT and P301S female mice (main effect of ethanol, F_1,35_ = 9.31, p = 0.004, Tukey’s multiple comparisons test, FWW vs FWE, p = 0.014) (Fig 2D).

**Figure 2.**
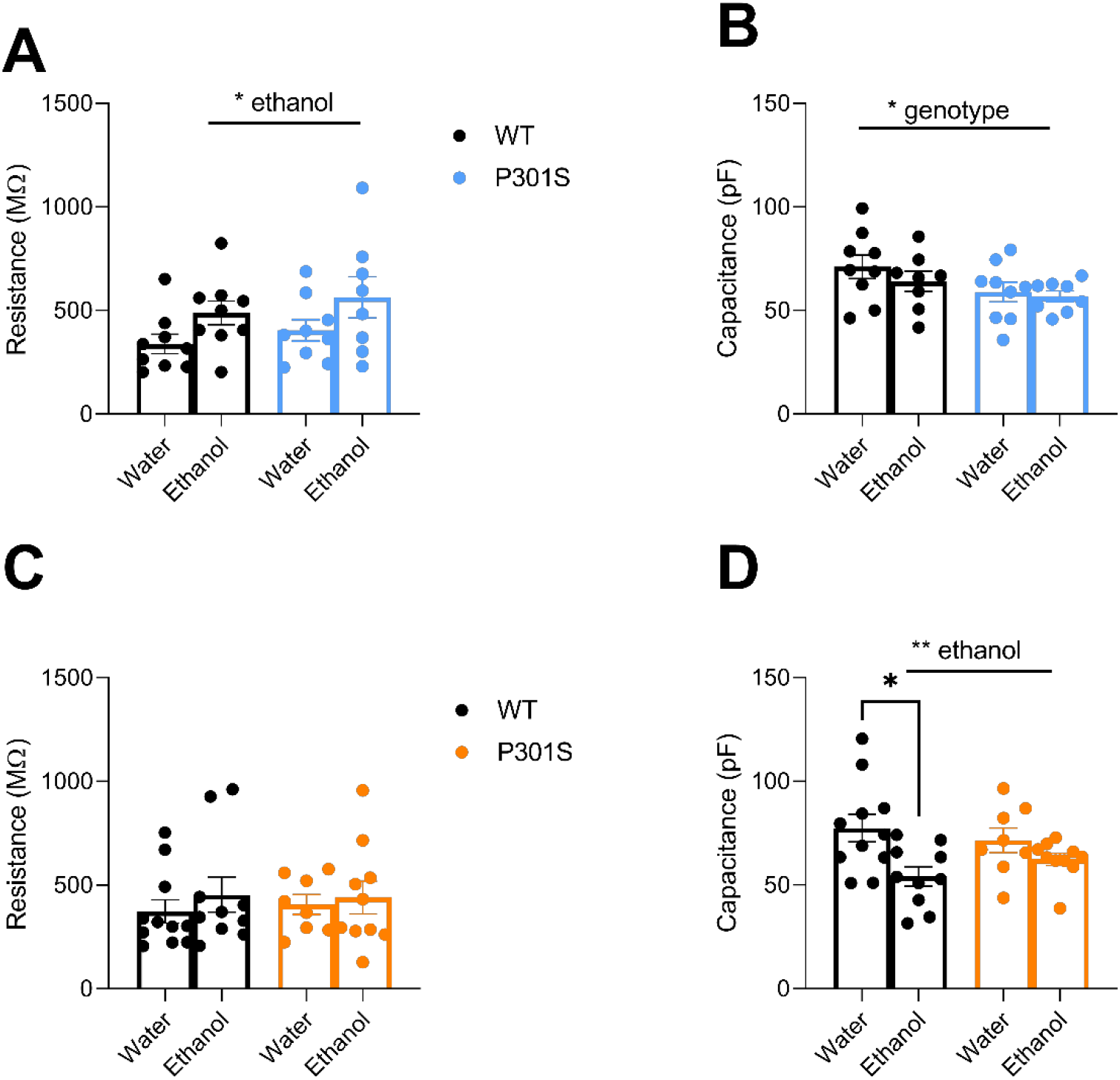
Electrophysiological properties of locus coeruleus (LC) neurons. (A) Input resistance is enhanced due to ethanol in male mice. (B) Capacitance of LC neurons is reduced in PS19 males. (C) Input resistance is unchanged in females. (C) Capacitance of LC neurons in reduced in ethanol-drinking females. Data expressed as mean ± S.E.M. The following sample sizes were used: MWW n = 4 mice, 8 cells, MWE n = 5 mice, 9 cells, MPW n = 5 mice, 9 cells, MPE n = 3 mice, 8 cells, FWW n = 4 mice, 11 cells, FWE n = 4 mice, 10 cells, FPW n = 4 mice, 8 cells, FPE n = 5 mice, 10 cells. * p ≤ 0.05. WT black bars (both sexes), P301S males blue bars, P301S females orange bars.

### Spontaneous excitatory postsynaptic currents

We next assessed the characteristics of spontaneous excitatory postsynaptic currents (sEPSCs) in the absence of tetrodotoxin. We found that male P301S animals had a significantly reduced frequency of sEPSCs compared to WT littermates (main effect of genotype, F_1,30_ = 15.95, p = 0.0004, Tukey’s multiple comparison test, p = 0.045) (Fig 3A for representative traces, Fig 3B). However, there were no significant changes observed due to ethanol consumption (main effect of ethanol, F_1,30_ = 0.52, p =0.48). There was a reduction in sEPSC amplitude in males due to ethanol consumption (main effect of ethanol, F_1,30_ = 4.89, p = 0.035) (Fig 3C). However, there were no significant differences due to genotype (main effect of genotype, F_1,30_ = 0.07, p = 0.80). We also observed a decrease in the decay time of sEPSCs in P301S males relative to WT littermates (main effect of genotype, F_1,30_ = 6.09, p = 0.20) (Fig 3D). There were no significant changes due to ethanol consumption (main effect of ethanol, F_1,30_ = 1.86, p = 0.18). While we observed a trend towards reduced sEPSC area in P301S males, we did not find any significant changes due to either genotype or ethanol consumption (main effect of genotype, F_1,30_ = 2.82, p = 0.10; main effect of ethanol, F_1,30_ = 0.02, p = 0.90) (Fig 3E).

**Figure 3.**
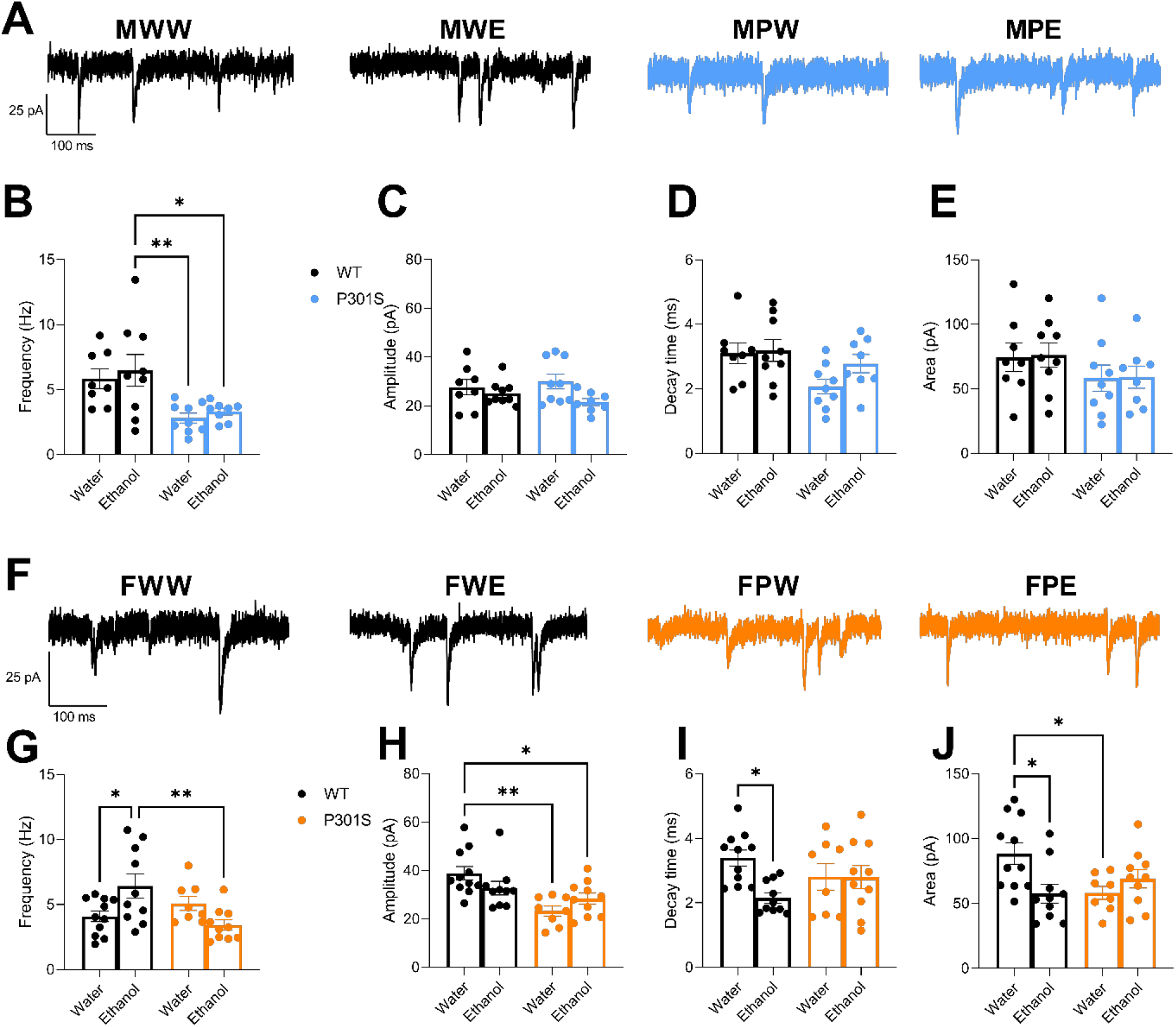
Spontaneous EPSCs in the LC are altered by ethanol consumption and P301S genotype. (A) Representative traces of sEPSCs from male LC neurons recorded in voltage clamp at −80 mV. (B) sEPSC frequency was reduced in P301S males. Main effect of genotype p < 0.01. (C) sEPSC amplitude was reduced by ethanol in males. Main effect of ethanol p < 0.05. (D) sEPSC decay time was reduced in P301S males. Main effect of genotype p < 0.05. (E) sEPSC area was unaltered in males. (F) Representative traces of sEPSCs from female LC neurons recorded in voltage clamp at −80 mV. (G) There was a significant interaction between P301S genotype and ethanol on sEPSC frequency in females. Interaction effect p < 0.01. (H) There was a significant interaction effect between P301S and ethanol on sEPSC amplitude in females and a significant reduction in amplitude in P301S female mice. Interaction effect p < 0.05, main effect of genotype p < 0.001. (I) There was a significant interaction effect between P301S and ethanol on sEPSC decay time in females and a significant reduction in decay time due to ethanol in WT females. Interaction effect p < 0.05, main effect of ethanol p < 0.05. (J) There was a significant interaction effect between P301S and ethanol on sEPSC area in females. Interaction effect p < 0.01. Data expressed as mean ± S.E.M. The following sample sizes were used: MWW n = 4 mice, 8 cells, MWE n = 5 mice, 9 cells, MPW n = 5 mice, 9 cells, MPE n = 3 mice, 8 cells, FWW n = 4 mice, 11 cells, FWE n = 4 mice, 10 cells, FPW n = 4 mice, 8 cells, FPE n = 5 mice, 10 cells. * p ≤ 0.05, ** p ≤ 0.01. WT black bars (both sexes), P301S males blue bars, P301S females orange bars.

We also assessed changes in sEPSCs in female animals separately (see Fig 3F for representative traces). We found that ethanol had opposite effects on the frequency of sEPSCs in WT and P301S mice. Ethanol enhanced sEPSC frequency in WT but decreased frequency in P301S animals (interaction effect, F_1,35_ = 10.46, p = 0.003, Tukey’s multiple comparisons test, FWW vs FWE, p = 0.039) (Fig 3G). However, we found an opposite effect on sEPSC amplitude. Ethanol decreased sEPSC amplitude in WT but modestly increased sEPSC amplitude in P301S mice (interaction effect, F_1,35_ = 4.51, p = 0.041). We also found significantly decreased sEPSC amplitude in P301S mice compared to WT controls (main effect of genotype, F_1,35_ = 14.55, p =0.0005, Tukey’s multiple comparisons tests, FWW vs FPW, p =0.001, FWW vs FPE, p = 0.026) (Fig 3H). We also found significant changes in the decay time of sEPSCs. Ethanol consumption induced a significant reduction in the decay time of sEPSCs in WT mice, but it had no effects in P301S mice (interaction effect, F_1,35_ = 4.31, p = 0.045; main effect of ethanol, F_1,35_ = 4.43, p = 0.043, Tukey’s multiple comparisons test, FWW vs FWE, p = 0.019) (Fig 3I). We also observed complimentary changes in sEPSC area. Ethanol consumption resulted in a significant decrease in sEPSC area in WT mice, but it had no effects in P301S animals (interaction effect, F_1,35_ = 7.94, p = 0.008, Tukey’s multiple comparisons test, FWW vs FWE, p = 0.02, FWW vs FPW, p = 0.036) (Fig 3J). Together these results demonstrate complex changes in excitatory inputs to the LC by ethanol and tau.

### Spontaneous action potentials

To investigate alterations in the spontaneous firing rate of LC neurons, we next assessed how P301S genotype and ethanol consumption impacted baseline action potential (AP) firing frequency and action potential properties of LC neurons. We found no significant changes in the baseline firing rate of LC neurons in male mice due to either genotype or ethanol consumption (main effect of ethanol, F_1,26_ = 1.65, p = 0.21; main effect of genotype, F_1,26_ = 0.55) (Fig 4A, B, C). However, we did find significant changes in the half-width of action potentials with opposing effects in WT and P301S mice males due to ethanol consumption. Ethanol increased AP half-width in WT but decreased AP half-width in P301S males (interaction effect, F_1,27_ = 10.92, p = 0.003, Tukey’s multiple comparisons test, FPW vs FPE, p = 0.015, FWE vs FPE, p = 0.017) (Fig 4D). Ethanol increased the magnitude of the after-hyperpolarization potential (AHP) in both WT and P301S males (main effect of ethanol, F_1,31_ = 35.52, p < 0.0001; Tukey’s multiple comparisons test, MWW vs MWE p = 0.009, MPW vs MPE p = 0.0001) (Fig 4E). Ethanol also reduced the AP threshold in male WT and P301S animals (main effect of ethanol, F_1,31_ = 38.18, p < 0.0001; Tukey’s multiple comparisons test, MWW vs MWE p < 0.0001, MPW vs MPE P = 0.0004) (Fig 4F). We saw no significant changes in either AP frequency (main effect of ethanol, F_1,34_ = 0.046, p = 0.49; main effect of genotype, F_1,36_ = 1.85, p = 0.18) (Fig 4G, H, I) or AP half-width (main effect of ethanol, F_1,34_ = 0.16, p = 0.69; main effect of genotype, F_1,34_ = 0.68, p =0.41) (Fig 4J) in female mice. Similar to males, ethanol enhanced the magnitude of (AHP) in both WT and P301S female mice (main effect of ethanol, F_1,35_ = 24.26, p < 0.0001; Tukey’s multiple comparisons test, FWW vs FWE p = 0.005, FPW vs FPE p = 0.009) (Fig 4K). Ethanol also reduced AP threshold in female WT and P301S animals (main effect of ethanol, F_1,35_ = 50.62, p < 0.0001; Tukey’s multiple comparisons test, FWW vs FWE p < 0.0001, FPW vs FPE p > 0.0001) (Fig 4L).

**Figure 4.**
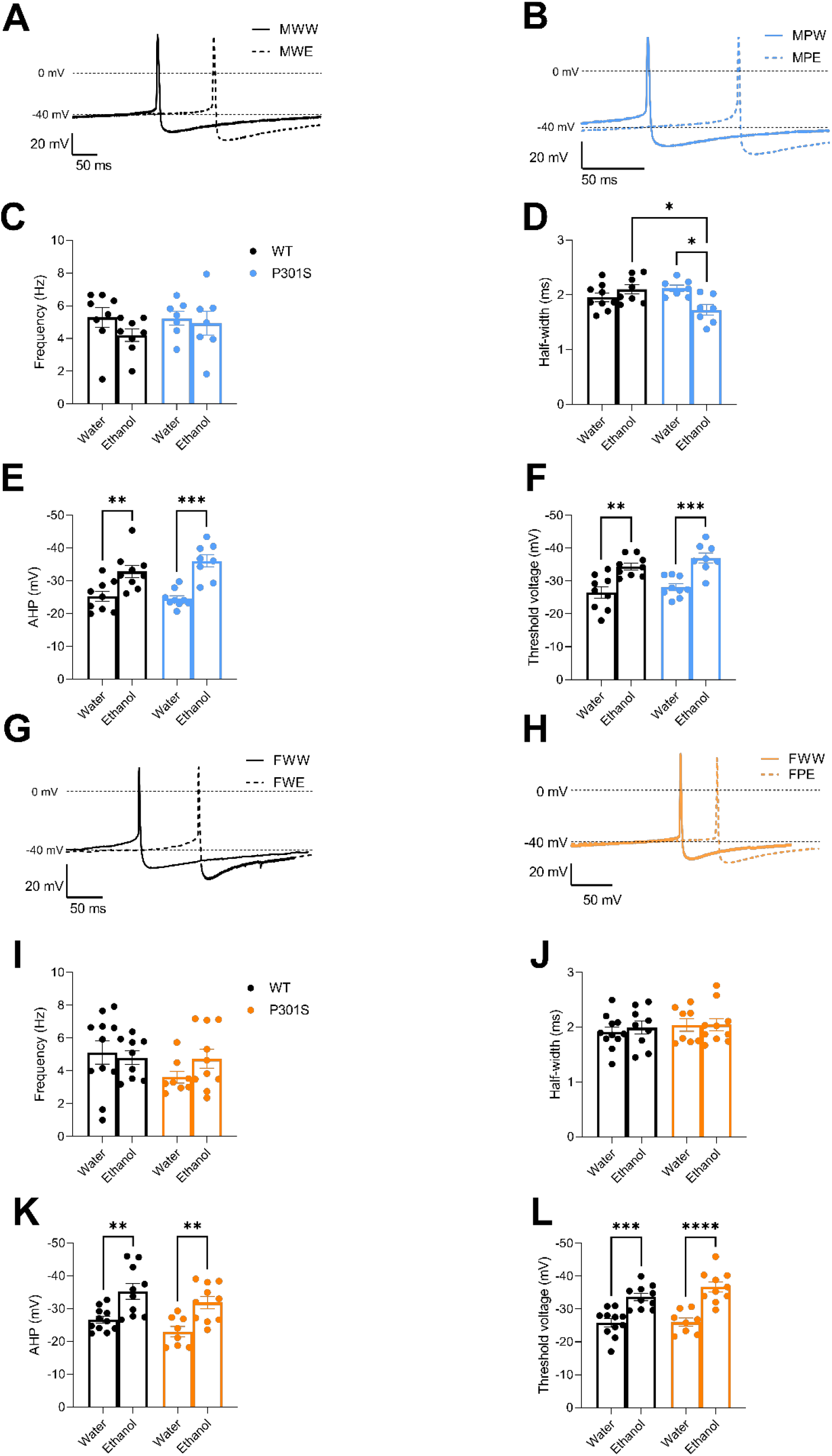
LC action potential kinetics are altered with no change in spontaneous firing rate. (A) Representative traces of MWW and MWE APs. (B) Representative traces of MPW and MPE APs. (C) AP frequency is unaltered in males. (D) There was a significant interaction effect between P301S and ethanol on AP half width in males. Interaction effect P < 0.01. (E) There was a larger AHP due to ethanol drinking in males. Main effect of ethanol p < 0.0001. (F) There was a lower threshold potential due to ethanol drinking in males. Main effect of ethanol p < 0.0001. (G) Representative traces of FWW and FWE APs. (H) Representative traces of FPW and FPE APs. (I) AP frequency is unaltered in females. (J) AP half width is unaltered in females. (K) There was a larger AHP due to ethanol drinking in females. Main effect of ethanol p < 0.0001. (L) There was a lower threshold potential due to ethanol drinking in females. Main effect of ethanol p < 0.0001. Data expressed as mean ± S.E.M. The following sample sizes were used: MWW n = 4 mice, 8 cells, MWE n = 5 mice, 8 cells, MPW n = 5 mice, 7 cells, MPE n = 3 mice, 7 cells, FWW n = 4 mice, 11 cells, FWE n = 4 mice, 9 cells, FPW n = 4 mice, 8 cells, FPE n = 5 mice, 10 cells. * p ≤ 0.05. WT black bars (both sexes), P301S males blue bars, P301S females orange bars.

### Excitability

We next assessed the excitability of LC neurons in response to both positive and negative current injections. We utilized negative current injection (0 to −100 pA, 20 pA per step) to measure input resistance of LC neurons in males (Fig 5A). In WT male mice, ethanol increased input resistance (input resistance x ethanol interaction effect, F_5,70_ = 16.09, p < 0.0001; main effect of ethanol, F_1,14_ = 20.76, p = 0.0004, Fig 5B). However, we did not observe the same increase in ethanol-induced input resistance in MPE versus MPW animals, as we saw in their WT littermates (main effect of ethanol, F_1,13_ = 0.313, p = 0.586, Fig 5C). We next measured the number of evoked action potentials fired in response to positive current injection (0 to 400 pA, 20 pA per step) in LC neurons, creating a firing curve in response to these graded current steps (Fig 5A). In male WT mice there were more action potentials fired after larger current injections. In MWE mice AP firing increased relative to current injection (firing curve x ethanol interaction effect, F_20, 280_ = 6.22, p < 0.0001, main effect of ethanol, F_1,14_ = 10.29, p = 0.006, Fig 5D). We observed a current injection by ethanol interaction in male P301S animals, however ethanol did not have a significant effect on its own (current step x ethanol interaction effect, F_20,260_ = 3.27, p < 0.001, main effect of ethanol, F_1,13_ = 4.30, p = 0.059, Fig 5E).

**Figure 5.**
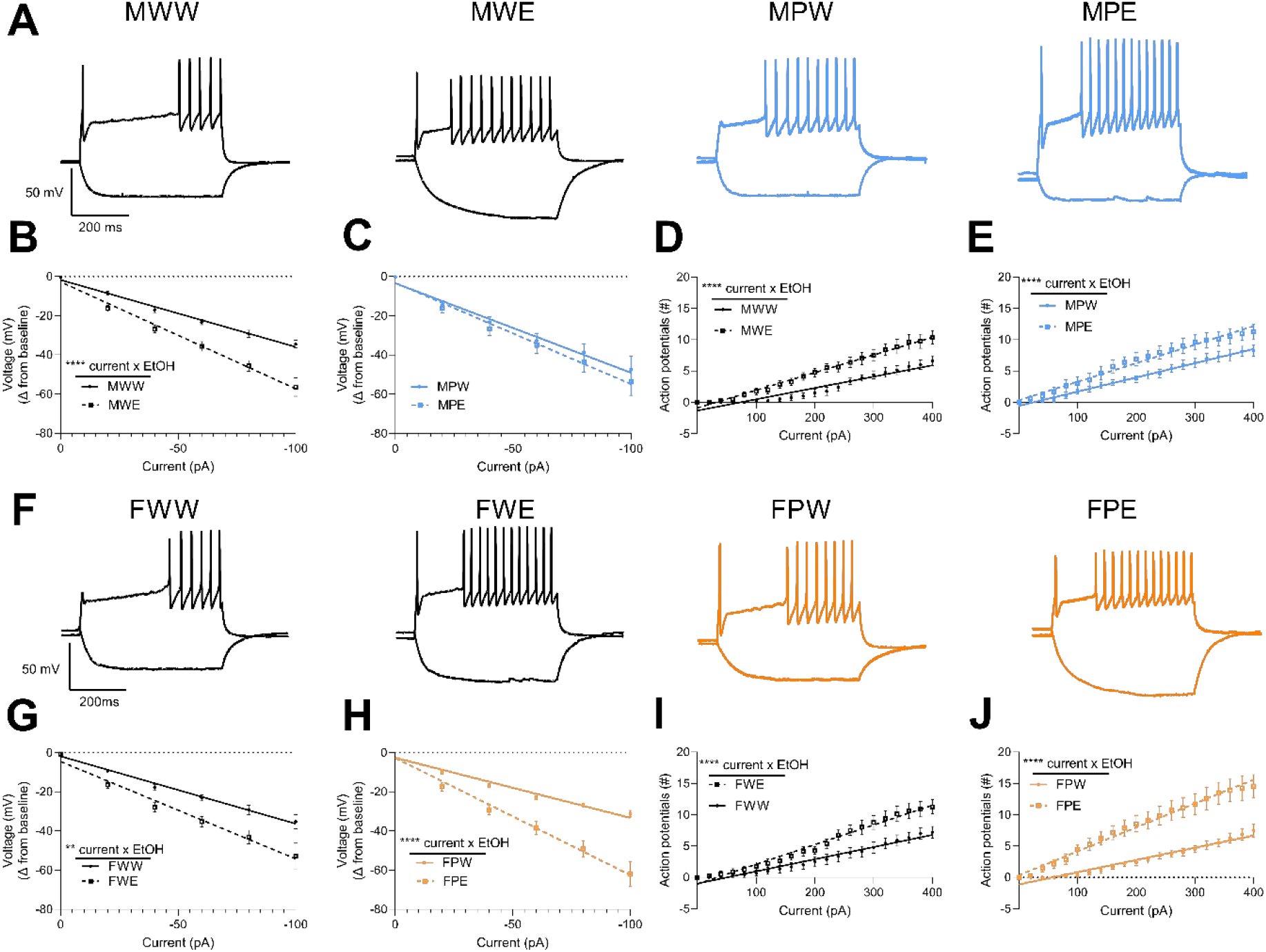
LC is hyperresponsive to current injections due to ethanol. (A) Representative data of LC response from male mice to −100 pA and 400 pA current injection while injecting a constant current to hold cells at −75 mV. (B) MWE mice have a greater input resistance than MWW. (C) No difference in hyperpolarization due to negative current injection between MPW and MPE mice. (D) LC neurons from MWE mice are more excitable than MWW mice. (E) LC neurons from MWE mice are more excitable than MWW mice. (F) Representative data of LC response from female mice to −100 pA and 400 pA current injection while injecting a constant current to hold cells at −75 mV. (G) FWE mice have a greater input resistance than FWW. (H) FPE mice have a greater input resistance than FPW. (I) LC neurons from FWE mice are more excitable than FWW mice. (J) LC neurons from FPE mice are more excitable than FPW mice. Data expressed as mean ± S.E.M. The following sample sizes were used: MWW n = 4 mice, 8 cells, MWE n = 5 mice, 8 cells, MPW n = 5 mice, 7 cells, MPE n = 3 mice, 8 cells, FWW n = 4 mice, 9 cells, FWE n = 4 mice, 8 cells, FPW n = 4 mice, 8 cells, FPE n = 5 mice, 7 cells. ** p ≤ 0.01, **** p ≤ 0.0001.

We performed the same experiment in female mice. In WT female mice, input resistance was also increased in ethanol consuming animals relative to controls (current step x ethanol interaction effect, F_5,75_ = 4.46, p = 0.001; main effect of ethanol, F_1,15_ = 12.28, p < 0.0001, Fig 5F & G). We observed a similar effect in P301S female mice where FPE had increased input resistance compared to FPW animals (input resistance x ethanol interaction effect, F_5,65_ = 23.13, p < 0.0001; main effect of ethanol, F_1,13_ = 25.84, p = 0.0002, Fig 5H). Interestingly, there was a trend towards an increase in input resistance in FPE and FPW vs FWE and FWW mice (3-way ANOVA, genotype x current x ethanol interaction, F_5,140_ = 2.21, p = 0.057). In female WT mice we observed increased evoked action potential firing with increased current injection in FWE versus FWW mice (firing curve x ethanol interaction effect, F_20,300_ = 6.09, p < 0.0001, main effect of drinking F_1,15_ = 6.51, p = 0.022, Fig 5I). We observed a similar ethanol-induced potentiation of LC action potential firing in female P301S animals (firing curve x ethanol interaction, F_20,260_ = 13.92, p < 0.0001, main effect of ethanol F_1,13_ = 19.10, p = 0.0008 (Fig 5J). Like our observations with hyperpolarizing current injection, we observed an increase in ethanol-induced potentiation of excitability in female P301S versus female WT animals (3-way ANOVA, current step x genotype x ethanol interaction effect, F_20,560_ = 1.86, p = 0.013). Taken together these results reveal a strong potentiation of LC excitability by long-term ethanol consumption.

## DISCUSSION

In this study, we examined drinking behavior and LC plasticity in aged (~7-month-old) male and female P301S mice and WT controls following 16 weeks of intermittent access to ethanol to assess the interaction between tauopathy and ethanol on LC physiology. Overall, total ethanol consumption did not differ between genotypes of either sex. We found that female mice drank more ethanol over the 16 weeks than male mice, regardless of genotype, while male P301S mice drank more and had a greater preference for ethanol over time than WT males. We further found complex effects of both P301S genotype and ethanol consumption on excitatory LC inputs and the kinetics of LC action potentials. We also found increased responsiveness of LC neurons to both excitatory and inhibitory currents in both male and female P301S and WT controls due to ethanol consumption.

Consistent with previous studies using the IA paradigm, we found that female mice drank more ethanol across the 16 weeks than male mice, irrespective of genotype (Hwa et al., 2011). When we examined the total amount of ethanol consumed each week, we found that male P301S mice drank more ethanol and had a greater preference for ethanol than WT controls during later weeks of the IA paradigm, suggesting that age-induced increase of tau accumulation in neurons may enhance ethanol consumption. However, we did not observe any genotype effects on weekly drinking in female mice. It is worth noting in our recent study that included more mice (Catavero et al., 2022), we similarly did not observe differences in total ethanol consumed by wild-type littermates or P301S mice, but we did observe differences in ethanol consumption over time in males, and ethanol preference in both male and female mice. While we did not assess tau pathology in this study, previous studies using the P301S mouse model have demonstrated that filamentous tau lesions begin to accumulate in the locus coeruleus and other brain regions of P310S mice at 6 months of age (Kang et al., 2020; Yoshiyama et al., 2007). Our P301S animals likely had tau lesions present during the latter part of the IA paradigm and at the 7-month time point used in our physiological studies. While we did not evaluate the effects of long-term ethanol consumption on tau accumulation, previous work using the triple transgenic AD mouse model (3xTg-AD), which includes a knock-in P301L tau variant along with additional AD-associated presenilin and amyloid precursor protein variants (Oddo et al., 2003), has demonstrated increased phosphorylated tau accumulation in the hippocampus, piriform cortex, medial prefrontal cortex, entorhinal cortex, and amygdala following both binge and long-term ethanol consumption (Barnett et al., 2022; Hoffman et al., 2019; Tucker et al., 2022). Future studies will need to investigate the impacts of alcohol consumption on tau accumulation in the LC itself. Hoffman and colleagues also found increased ethanol consumption at the start of a 2-bottle choice paradigm in 3xTg-AD mice versus WT controls (Hoffman et al., 2019). However, our study observed increased ethanol consumption during later time points of our paradigm, this could reflect differences in potentiation of ethanol drinking caused by abstinence in our IA paradigm or due to the onset of tau tangles at the ~6-month time point.

While our IA paradigm revealed differences in alcohol consumption and preference in males based on genotype, it limited direct sex-based comparisons in our physiological studies due to increased ethanol consumption in females. Accordingly, we analyzed our physiological data separately for male and female animals. In female WT animals that consumed ethanol, we observed an increase in sEPSC frequency, along with a decrease in total current transfer (area) due to decreased amplitude and decay kinetics. These findings are suggestive of an increased insertion of Ca^2+^-permeable AMPA receptors (CP-AMPAR) (Cull-Candy et al., 2006). Increased CP-AMPAR insertion is known to promote long-term potentiation in other neuronal populations (Guire et al., 2008; Park et al., 2018). Previous studies have implicated increased CP-AMPAR insertion in neurons in the basolateral amygdala and ventral tegmental area in mouse models of alcohol consumption, and our results suggest similar changes in LC (Faccidomo et al., 2021; Hausknecht et al., 2015; Kircher et al., 2019). Curiously, we found opposite effects of ethanol consumption on sEPSC frequency and amplitude and no effect on sEPSC decay kinetics due to ethanol consumption in P301S females compared to WT females. However, there is evidence to suggest that the accumulation of pathological, phosphorylated tau disrupts normal post-synaptic density structure and specifically disrupts and reduces AMPA receptor insertion on the synapse (Gong and Lippa, 2010; Hoover et al., 2010; Jurado, 2017; Shrivastava et al., 2019). Disruptions in AMPA receptor trafficking could disrupt this ethanol-induced plasticity in P301S females. Interestingly, compared to water drinking animals alone, P301S females exhibited significantly reduced sEPSC amplitude and charge transfer, reflecting a loss of excitatory synaptic drive on LC neurons specifically in female animals.

In males, we observed more changes in excitatory inputs due to the P301S genotype than due to IA to ethanol. This could suggest an effect of dose, in that males did not drink sufficient ethanol to modulate excitatory LC synapses. Critically, we did not observe the same increase in sEPSC frequency and decrease in kinetics that we observed in female WT mice in the male WT subjects. This may reflect decreased ethanol consumption in males, or it may reflect sex differences in LC plasticity in response to long-term ethanol consumption. While sex differences in LC plasticity have not been studied specifically in the context of alcohol use, structural and molecular differences in the LC based on sex are well established (Bangasser et al., 2011; Mulvey et al., 2018), as well as sexually dimorphic responses to other behavioral phenomena including stress (Curtis et al., 2006). We did find decreased frequency of sEPSC in P301S males in both water and ethanol groups, which could reflect a decrease in excitatory synapse number in the LC as discussed with the female P301S animals or a decrease in glutamate release. Changes in AMPA receptor subunit trafficking due to tau accumulation could also explain the decrease in decay kinetics we observed in male P301S mice (Hoover et al., 2010).

While we did not observe baseline changes in LC spontaneous firing rates due to IA to ethanol or P301S genotype, we did observe significant changes in excitability of these neurons. In both male and female P301S and WT controls we found a significant increase in excitability due to ethanol consumption, which agrees with previous anesthetized *in vivo* recordings demonstrating increased excitability of the LC following alcohol withdrawal and abstinence (Engberg and Hajós, 1992). Along with increased firing we saw an increase in AHP and a hyperpolarized AP threshold in ethanol-consuming mice of all genotypes and sexes. The increased firing rate may reflect ethanol-dependent adaptation in the function of voltage-gated Ca^2+^ channel function and small-conductance Ca^2+^-activated K^+^ (SK) channels. Indeed, a decrease in SK channel function has been previously observed in rodent alcohol consumption in nucleus accumbens medium spiny neurons and hippocampal pyramidal neurons respectively (Shan et al., 2019; Uhrig et al., 2017). Furthermore, ethanol experience reduces SK channel currents in the lateral orbital frontal cortex (Nimitvilai et al., 2016), and disrupts interactions between SK channels and glutamatergic synapses in the hippocampus (Mulholland et al., 2011). In the ventral tegmental area, ethanol reduces the function of SK channels, which may enhance their burst firing modes (Mulholland et al., 2009). Given that SK2 channels are the most prevalent form in the LC and SK2 channel reduction promotes faster firing rates (Matschke et al., 2018), future studies could investigate whether SK2 channel function is reduced in the LC following ethanol administration. While reduced SK channel expression would be hypothesized to reduce AHP, increased Ca^2+^ influx may drive a greater SK channel mediated AHP with reduced total SK channel numbers (Faber and Sah, 2002). Future studies could assess Ca^2+^dynamics or L-type Ca^2+^ channel function to further investigate the cause of the increased AHP along with increased excitability. Further, we found that female P301S mice had a greater magnitude of increase in excitability due to ethanol. Current data on the effects of tau pathology on neuronal excitability are conflicting, with some studies finding increased or decreased excitability in cortical or hippocampal neurons (Busche et al., 2019; Crimins et al., 2012; Huijbers et al., 2019). While the mechanism behind these changes is unclear, tau may disrupt a variety of voltage-gated ion channels. Tau reduces Kv4.2 in dendrites of CA1 neurons in the hippocampus (Hall et al., 2015). Future studies should further characterize the effects of tau accumulation on ion channel conductance in the LC.

Over the past decade it is becoming more apparent that the LC is not a homogenous nucleus, but has modular functionality based on outputs to different nuclei (Chandler et al., 2019). In particular, electrophysiological signatures have been observed for multiple outputs, and these may be liable to plasticity induced by experience (stressors) (Borodovitsyna et al., 2020; Chandler et al., 2014; Li et al., 2016). Moreover, there appear to be discrete modules that influence different motivated behaviors (Breton-Provencher et al., 2022; Hickey et al., 2014; Hirschberg et al., 2017). Future studies investigating how tau pathology and ethanol alter these individual modules would be undoubtedly insightful.

Taken together our data demonstrate significant changes in both excitatory inputs to the LC and intrinsic LC excitability due to tauopathy and long-term ethanol consumption. Given the broad role of the LC in diverse neurological process, increased excitability of the LC could contribute to the development of anxiety-like phenotypes, learning and memory deficits, and alterations to arousal, all of which are known to be altered in AD, FTD, and alcohol use disorder (Cerejeira et al., 2012). Future studies will be critical to further our understanding of behavioral consequences of altered LC function in the context of both alcohol use and tauopathy.

## ACKNOWLEDGMENTS

This work was funded by U01AA020911-0851 (ZAM), R01DA049261 (ZAM), and T32AA007573 (AMD).

## LITERATURE CITED

Andrés-Benito, P., et al., 2017. Locus coeruleus at asymptomatic early and middle Braak stages of neurofibrillary tangle pathology. Neuropathol Appl Neurobiol. 43, 373–392.

Aston-Jones, G., Cohen, J. D., 2005. An integrative theory of locus coeruleus-norepinephrine function: adaptive gain and optimal performance. Annu Rev Neurosci. 28, 403–50.

Bailly, D., et al., 1992. Effects of beta-blocking drugs in alcohol withdrawal: a double-blind comparative study with propranolol and diazepam. Biomed Pharmacother. 46, 419–24.

Bangasser, D. A., et al., 2011. Sexual dimorphism in locus coeruleus dendritic morphology: a structural basis for sex differences in emotional arousal. Physiol Behav. 103, 342–51.

Barnett, A., et al., 2022. Adolescent Binge Alcohol Enhances Early Alzheimer’s Disease Pathology in Adulthood Through Proinflammatory Neuroimmune Activation. Front Pharmacol. 13, 884170.

Beardmore, R., et al., 2021. The Locus Coeruleus in Aging and Alzheimer’s Disease: A Postmortem and Brain Imaging Review. J Alzheimers Dis. 83, 5–22.

Bondareff, W., et al., 1982. Loss of neurons of origin of the adrenergic projection to cerebral cortex (nucleus locus ceruleus) in senile dementia. Neurology. 32, 164–8.

Borodovitsyna, O., et al., 2020. Anatomically and functionally distinct locus coeruleus efferents mediate opposing effects on anxiety-like behavior. Neurobiol Stress. 13, 100284.

Braak, H., Braak, E., 1991. Neuropathological stageing of Alzheimer-related changes. Acta Neuropathol. 82, 239–59.

Breslow, R. A., et al., 2017. Trends in Alcohol Consumption Among Older Americans: National Health Interview Surveys, 1997 to 2014. Alcohol Clin Exp Res. 41, 976–986.

Breton-Provencher, V., et al., 2022. Spatiotemporal dynamics of noradrenaline during learned behaviour. Nature. 606, 732–738.

Burnham, N. W., Thiele, T. E., 2017. Voluntary Binge-like Ethanol Consumption Site-specifically Increases c-Fos Immunoexpression in Male C57BL6/J Mice. Neuroscience. 367, 159–168.

Busche, M. A., et al., 2019. Tau impairs neural circuits, dominating amyloid-β effects, in Alzheimer models in vivo. Nat Neurosci. 22, 57–64.

Cannady, R., et al., 2018. Chronic Alcohol, Intrinsic Excitability, and Potassium Channels: Neuroadaptations and Drinking Behavior. Handb Exp Pharmacol. 248, 311–343.

Catavero, C. M., et al., 2022. Effects of Long-Term Alcohol Consumption on Behavior in the P301S (Line PS19) Tauopathy Mouse Model. bioRxiv. 2022.07.12.499737.

Cerejeira, J., et al., 2012. Behavioral and Psychological Symptoms of Dementia. Frontiers in Neurology. 3.

Chalermpalanupap, T., et al., 2018. Locus Coeruleus Ablation Exacerbates Cognitive Deficits, Neuropathology, and Lethality in P301S Tau Transgenic Mice. J Neurosci. 38, 74–92.

Chandler, D. J., et al., 2014. Heterogeneous organization of the locus coeruleus projections to prefrontal and motor cortices. Proc Natl Acad Sci U S A. 111, 6816–21.

Chandler, D. J., et al., 2019. Redefining Noradrenergic Neuromodulation of Behavior: Impacts of a Modular Locus Coeruleus Architecture. J Neurosci. 39, 8239–8249.

Ciampa, C. J., et al., 2022. Associations among locus coeruleus catecholamines, tau pathology, and memory in aging. Neuropsychopharmacology. 47, 1106–1113.

Crimins, J. L., et al., 2012. Electrophysiological changes precede morphological changes to frontal cortical pyramidal neurons in the rTg4510 mouse model of progressive tauopathy. Acta Neuropathol. 124, 777–95.

Cull-Candy, S., et al., 2006. Regulation of Ca2+-permeable AMPA receptors: synaptic plasticity and beyond. Curr Opin Neurobiol. 16, 288–97.

Curtis, A. L., et al., 2006. Sexually Dimorphic Responses of the Brain Norepinephrine System to Stress and Corticotropin-Releasing Factor. Neuropsychopharmacology. 31, 544–554.

Downs, A. M., McElligott, Z. A., 2022. Noradrenergic circuits and signaling in substance use disorders. Neuropharmacology. 208, 108997.

Engberg, G., Hajós, M., 1992. Alcohol withdrawal reaction as a result of adaptive changes of excitatory amino acid receptors. Naunyn Schmiedebergs Arch Pharmacol. 346, 437–41.

Faber, E. S., Sah, P., 2002. Physiological role of calcium-activated potassium currents in the rat lateral amygdala. J Neurosci. 22, 1618–28.

Faccidomo, S., et al., 2021. Calcium-permeable AMPA receptor activity and GluA1 trafficking in the basolateral amygdala regulate operant alcohol self-administration. Addict Biol. 26, e13049.

Funk, C. K., et al., 2006. Corticotropin-releasing factor within the central nucleus of the amygdala mediates enhanced ethanol self-administration in withdrawn, ethanol-dependent rats. J Neurosci. 26, 11324–32.

Gilpin, N. W., et al., 2015. The central amygdala as an integrative hub for anxiety and alcohol use disorders. Biol Psychiatry. 77, 859–69.

Gong, Y., Lippa, C. F., 2010. Review: disruption of the postsynaptic density in Alzheimer’s disease and other neurodegenerative dementias. Am J Alzheimers Dis Other Demen. 25, 547–55.

Gottlieb, L. D., et al., 1994. Randomized controlled trial in alcohol relapse prevention: role of atenolol, alcohol craving, and treatment adherence. J Subst Abuse Treat. 11, 253–8.

Grucza, R. A., et al., 2018. Trends in Adult Alcohol Use and Binge Drinking in the Early 21st-Century United States: A Meta-Analysis of 6 National Survey Series. Alcohol Clin Exp Res. 42, 1939–1950.

Guire, E. S., et al., 2008. Recruitment of calcium-permeable AMPA receptors during synaptic potentiation is regulated by CaM-kinase I. J Neurosci. 28, 6000–9.

Haass-Koffler, C. L., et al., 2017. Higher pretreatment blood pressure is associated with greater alcohol drinking reduction in alcohol-dependent individuals treated with doxazosin. Drug Alcohol Depend. 177, 23–28.

Hall, A. M., et al., 2015. Tau-Dependent Kv4.2 Depletion and Dendritic Hyperexcitability in a Mouse Model of Alzheimer’s Disease. The Journal of Neuroscience. 35, 6221–6230.

Harley, C. W., 1987. A role for norepinephrine in arousal, emotion and learning?: limbic modulation by norepinephrine and the Kety hypothesis. Prog Neuropsychopharmacol Biol Psychiatry. 11, 419–58.

Harrison, N. L., et al., 2017. Effects of acute alcohol on excitability in the CNS. Neuropharmacology. 122, 36–45.

Hausknecht, K., et al., 2015. Excitatory synaptic function and plasticity is persistently altered in ventral tegmental area dopamine neurons after prenatal ethanol exposure. Neuropsychopharmacology. 40, 893–905.

Heymann, D., et al., 2016. The Association Between Alcohol Use and the Progression of Alzheimer’s Disease. Curr Alzheimer Res. 13, 1356–1362.

Hickey, L., et al., 2014. Optoactivation of locus ceruleus neurons evokes bidirectional changes in thermal nociception in rats. J Neurosci. 34, 4148–60.

Hirschberg, S., et al., 2017. Functional dichotomy in spinal-vs prefrontal-projecting locus coeruleus modules splits descending noradrenergic analgesia from ascending aversion and anxiety in rats. Elife. 6.

Hoffman, J. L., et al., 2019. Alcohol drinking exacerbates neural and behavioral pathology in the 3xTg-AD mouse model of Alzheimer’s disease. Int Rev Neurobiol. 148, 169–230.

Hoover, B. R., et al., 2010. Tau mislocalization to dendritic spines mediates synaptic dysfunction independently of neurodegeneration. Neuron. 68, 1067–81.

Horwitz, R. I., et al., 1989. The efficacy of atenolol in the outpatient management of the alcohol withdrawal syndrome. Results of a randomized clinical trial. Arch Intern Med. 149, 1089–93.

Huijbers, W., et al., 2019. Tau Accumulation in Clinically Normal Older Adults Is Associated with Hippocampal Hyperactivity. J Neurosci. 39, 548–556.

Hwa, L. S., et al., 2011. Persistent escalation of alcohol drinking in C57BL/6J mice with intermittent access to 20% ethanol. Alcohol Clin Exp Res. 35, 1938–47.

Iba, M., et al., 2015. Tau pathology spread in PS19 tau transgenic mice following locus coeruleus (LC) injections of synthetic tau fibrils is determined by the LC’s afferent and efferent connections. Acta Neuropathol. 130, 349–62.

Jacobs, H. I. L., et al., 2021. In vivo and neuropathology data support locus coeruleus integrity as indicator of Alzheimer’s disease pathology and cognitive decline. Sci Transl Med. 13,eabj2511.

Jurado, S., 2017. AMPA Receptor Trafficking in Natural and Pathological Aging. Front Mol Neurosci. 10, 446.

Kang, S. S., et al., 2020. Norepinephrine metabolite DOPEGAL activates AEP and pathological Tau aggregation in locus coeruleus. J Clin Invest. 130, 422–437.

Kayama, Y., Koyama, Y., 2003. Control of sleep and wakefulness by brainstem monoaminergic and cholinergic neurons. Acta Neurochir Suppl. 87, 3–6.

Keaney, F., et al., 2001. A double-blind randomized placebo-controlled trial of lofexidine in alcohol withdrawal: lofexidine is not a useful adjunct to chlordiazepoxide. Alcohol Alcohol. 36, 426–30.

Kircher, D. M., et al., 2019. Ethanol Experience Enhances Glutamatergic Ventral Hippocampal Inputs to D1 Receptor-Expressing Medium Spiny Neurons in the Nucleus Accumbens Shell. J Neurosci. 39, 2459–2469.

Koob, G. F., 2015. The dark side of emotion: the addiction perspective. Eur J Pharmacol. 753, 73–87.

Li, Y., et al., 2016. Retrograde optogenetic characterization of the pontospinal module of the locus coeruleus with a canine adenoviral vector. Brain Res. 1641, 274–90.

Luster, B. R., et al., 2020. Inhibitory transmission in the bed nucleus of the stria terminalis in male and female mice following morphine withdrawal. Addict Biol. 25, e12748.

Mason, S. T., 1981. Noradrenaline in the brain: progress in theories of behavioural function. Prog Neurobiol. 16, 263–303.

Masters, C. L., et al., 2015. Alzheimer’s disease. Nat Rev Dis Primers. 1, 15056.

Matschke, L. A., et al., 2018. Calcium-activated SK potassium channels are key modulators of the pacemaker frequency in locus coeruleus neurons. Molecular and Cellular Neuroscience. 88, 330–341.

Mayeux, R., Stern, Y., 2012. Epidemiology of Alzheimer disease. Cold Spring Harb Perspect Med. 2.

Milivojevic, V., et al., 2020. Effects of Prazosin on Provoked Alcohol Craving and Autonomic and Neuroendocrine Response to Stress in Alcohol Use Disorder. Alcohol Clin Exp Res. 44, 1488–1496.

Mulholland, P. J., et al., 2011. Small conductance calcium-activated potassium type 2 channels regulate alcohol-associated plasticity of glutamatergic synapses. Biol Psychiatry. 69, 625–32.

Mulholland, P. J., et al., 2009. Sizing up ethanol-induced plasticity: the role of small and large conductance calcium-activated potassium channels. Alcohol Clin Exp Res. 33, 1125–35.

Mulvey, B., et al., 2018. Molecular and Functional Sex Differences of Noradrenergic Neurons in the Mouse Locus Coeruleus. Cell Reports. 23, 2225–2235.

Nimitvilai, S., et al., 2016. Chronic Intermittent Ethanol Exposure Enhances the Excitability and Synaptic Plasticity of Lateral Orbitofrontal Cortex Neurons and Induces a Tolerance to the Acute Inhibitory Actions of Ethanol. Neuropsychopharmacology. 41, 1112–27.

Oddo, S., et al., 2003. Triple-transgenic model of Alzheimer’s disease with plaques and tangles: intracellular Abeta and synaptic dysfunction. Neuron. 39, 409–21.

Ohm, D. T., et al., 2020. Degeneration of the locus coeruleus is a common feature of tauopathies and distinct from TDP-43 proteinopathies in the frontotemporal lobar degeneration spectrum. Acta Neuropathol. 140, 675–693.

Paladini, C. A., et al., 2007. Electrophysiological properties of catecholaminergic neurons in the norepinephrine-deficient mouse. Neuroscience. 144, 1067–74.

Park, P., et al., 2018. The Role of Calcium-Permeable AMPARs in Long-Term Potentiation at Principal Neurons in the Rodent Hippocampus. Front Synaptic Neurosci. 10, 42.

Peng, B., et al., 2020. Role of Alcohol Drinking in Alzheimer’s Disease, Parkinson’s Disease, and Amyotrophic Lateral Sclerosis. Int J Mol Sci. 21.

Poe, G. R., et al., 2020. Locus coeruleus: a new look at the blue spot. Nature Reviews Neuroscience. 21, 644–659.

Rehm, J., et al., 2019. Alcohol use and dementia: a systematic scoping review. Alzheimers Res Ther. 11, 1.

Rorabaugh, J. M., et al., 2017. Chemogenetic locus coeruleus activation restores reversal learning in a rat model of Alzheimer’s disease. Brain. 140, 3023–3038.

Rüb, U., et al., 2001. The autonomic higher order processing nuclei of the lower brain stem are among the early targets of the Alzheimer’s disease-related cytoskeletal pathology. Acta Neuropathol. 101, 555–64.

Shan, L., et al., 2019. Nucleus accumbens shell small conductance potassium channels underlie adolescent ethanol exposure-induced anxiety. Neuropsychopharmacology. 44, 1886–1895.

Shrivastava, A. N., et al., 2019. Clustering of Tau fibrils impairs the synaptic composition of α3-Na(+)/K(+)-ATPase and AMPA receptors. Embo j. 38.

Simpson, T. L., et al., 2015. A pilot trial of prazosin, an alpha-1 adrenergic antagonist, for comorbid alcohol dependence and posttraumatic stress disorder. Alcohol Clin Exp Res. 39, 808–17.

Simpson, T. L., et al., 2018. Double-Blind Randomized Clinical Trial of Prazosin for Alcohol Use Disorder. Am J Psychiatry. 175, 1216–1224.

Sinha, R., et al., 2021. Moderation of Prazosin’s Efficacy by Alcohol Withdrawal Symptoms. Am J Psychiatry. 178, 447–458.

Thiele, T. E., et al., 2000. Ethanol-induced c-fos expression in catecholamine-and neuropeptide Y-producing neurons in rat brainstem. Alcohol Clin Exp Res. 24, 802–9.

Tucker, A. E., et al., 2022. Chronic Ethanol Causes Persistent Increases in Alzheimer’s Tau Pathology in Female 3xTg-AD Mice: A Potential Role for Lysosomal Impairment. Front Behav Neurosci. 16, 886634.

Tyas, S. L., 2001. Alcohol use and the risk of developing Alzheimer’s disease. Alcohol Res Health. 25, 299–306.

Uhrig, S., et al., 2017. Differential Roles for L-Type Calcium Channel Subtypes in Alcohol Dependence. Neuropsychopharmacology. 42, 1058–1069.

Vazey, E. M., et al., 2018. Central Noradrenergic Interactions with Alcohol and Regulation of Alcohol-Related Behaviors. Handb Exp Pharmacol. 248, 239–260.

Wang, Z. T., et al., 2021. Associations of Alcohol Consumption with Cerebrospinal Fluid Biomarkers of Alzheimer’s Disease Pathology in Cognitively Intact Older Adults: The CABLE Study. J Alzheimers Dis. 82, 1045–1054.

Weinshenker, D., et al., 2000. Ethanol-associated behaviors of mice lacking norepinephrine. J Neurosci. 20, 3157–64.

Williams, J. T., et al., 1984. Membrane properties of rat locus coeruleus neurones. Neuroscience. 13, 137–56.

Wilson, S. R., et al., 2014. The prevalence of harmful and hazardous alcohol consumption in older U.S. adults: data from the 2005-2008 National Health and Nutrition Examination Survey (NHANES). J Gen Intern Med. 29, 312–9.

Yoshiyama, Y., et al., 2007. Synapse loss and microglial activation precede tangles in a P301S tauopathy mouse model. Neuron. 53, 337–51.

Zweig, R. M., et al., 1988. The neuropathology of aminergic nuclei in Alzheimer’s disease. Ann Neurol. 24, 233–42.

